# Ontology-aware DNA methylation classification with a curated atlas of human tissues and cell types

**DOI:** 10.1101/2025.04.18.649618

**Authors:** Mirae Kim, Ruth Dannenfelser, Yufei Cui, Genevera Allen, Vicky Yao

## Abstract

DNA methylation (DNAm) is a core gene regulatory mechanism that captures cellular responses to short- and long-term stimuli such as environmental exposures, aging, and cellular differentiation. Although DNAm has proven valuable as a baseline biomarker for aging by enabling robust characterization of disease-associated methylation shifts associated with age, its potential to reveal analogous shifts in the context of tissue remains underexplored. A major obstacle has been the absence of comprehensive, curated reference atlases spanning diverse normal human tissues, limiting most existing work to disease-subtype differentiation or localized tissue comparisons. To bridge this gap, we assemble the largest and most diverse atlas of exclusively healthy human tissue and cell samples profiled by 450K arrays, comprising of 16,959 samples across 86 tissues and cell types. Leveraging this resource, we introduce an ontology-aware classification framework that identifies robust CpG features associated with tissue and cell identity while integrating known anatomical and functional relationships (e.g., prefrontal cortex in the brain, leukocytes in blood). Our novel application of Minipatch learning distills a set of 190 CpG sites that can accurately support multi-label classification. We further validate our approach through an ontology-based label transfer task, demonstrating the effectiveness of ontology-informed learning to accurately predict relevant labels for 31 tissues and cell types not seen during training. These findings underscore the potential of our framework to enhance our understanding of healthy methylation landscapes and facilitate future applications in disease detection and personalized medicine.

## Introduction

DNA methylation (DNAm) is a key epigenetic mechanism responsible for regulating gene expression and chromatin organization, serving both to preserve cell lineage identity and dynamically mediate cellular responses to environmental stimuli [1]. Due to these dual roles, DNAm patterns have emerged as powerful biomarkers for a variety of biological processes, including tissue and cell-type classification based on conserved methylation signatures [2, 3], quantification of aging-associated methylation shifts [4, 5], and detection of molecular alterations associated with disease [6, 7] or environmental exposures [8, 9]. These broad detection abilities have led to a flurry of excitement about the potential of DNAm as a comprehensive molecular snapshot of human health, with tissue and cell type classification methods as central components of this vision [10, 11].

Existing tissue and cell classification approaches typically aim to identify stable sets of unique, distinguishing methylation patterns. These patterns are often localized to unmethylated regulatory regions that are important in defining cellular identity and function [1]. Because cell lineage is a primary driver of these conserved methylation signatures, considerable effort has been devoted to computational deconvolution methods that infer cell-type proportions from bulk-tissue methylation profiles [10, 12]. Despite advances in reference-free deconvolution approaches, identification of reliable tissue- and cell-type markers still rely heavily on statistical analyses of DNA methylation reference datasets [13].

The effectiveness of these methods is closely tied to the comprehensiveness and expressiveness of their underlying reference datasets. Ideally, reference datasets would capture extensive tissue and cell type diversity with broad coverage of methylation sites across the genome. In practice, researchers must balance the tradeoff between array-based technologies (e.g., 450K, EPIC arrays), offering broad sample diversity at lower genomic coverage, and sequencing-based technologies (e.g., whole genome bisulfite sequencing (WGBS), reduced representation bisulfite sequencing (RRBS)), which provide greater genomic coverage across CpG sites but at higher costs and lower sample throughput. Recent advances in single-cell DNA methylation techniques promise unprecedented cellular resolution [14, 15] but, their high cost, limited scalability, and intrinsic data sparsity currently limit their use for large-scale tissue profiling [16, 17]. Consequently, comprehensive bulk-tissue methylation reference collections remain essential. Various DNAm tissue and cell-type atlases have been assembled across arrays [18, 19], RRBS [20–22], and WGBS [2, 3]. However, some of these efforts combine both healthy and diseased samples to achieve broader tissue coverage, but can inadvertently obscure signals specific to healthy cellular states [23]. Meanwhile, even the most comprehensive healthy atlas, profiled using WGBS, covers only 39 cell types from 18 major tissues [3], missing entire organ systems (e.g., the male reproductive system) and is limited to a few representative cell types per tissue.

Here, we address these gaps by assembling, to our knowledge, the largest curated atlas of exclusively healthy, primary human tissue and cell types. Our data compendium spans 55 tissue and cell types, sourced from 210 publicly available studies profiled by 450K arrays within the Gene Expression Ominbus (GEO) [24]. Although 450K arrays profile fewer CpG sites than WGBS, they still robustly capture cell-type specific methylation signals [25], and their widespread usage has resulted in unmatched sample diversity. Leveraging this comprehensive atlas, we introduce a multi-label, ontology aware classification framework explicitly designed to prioritize tissue identity rather than cell type composition. Unlike traditional deconvolution methods, our ontology-driven approach incorporates known functional and lineage relationships between tissues and cell types, enabling identification of CpG sites that represent distinct biological signatures beyond lineage origin alone. By anchoring our approach within a structured anatomical and functional ontology, we can also effectively model intermediate nodes and organ-system-level relationships, ultimately expanding our classification coverage to 72 distinct anatomical entities. Lastly, by leveraging the lineage and functional relationships encoded by the ontology, our model achieves accurate predictions of related labels for 31 tissues and cell types unseen during training, demonstrating robust and biologically meaningful generalizability despite inherent gaps in available reference data.

## Results

### Curating a diverse DNA methylation compendium of 86 healthy primary tissues and cell types

We assembled a reference compendium of DNA methylation (DNAm) profiles across a wide variety of healthy, untreated, primary tissues and cell types to model DNAm relationships in the context of anatomical and functional similarity. To this end, we obtained all publicly available Illumina 450K data deposited in Gene Expression Omnibus (GEO) [24] and manually filtered out samples that were diseased, treated, or derived from cell lines or organoids. We then reprocessed all files using a consistent data processing pipeline, removing samples that did not pass quality control checks (see Methods). For consistency, we disambiguated sample type labels by manually reconciling them with the UBERON Tissue Ontology [26], yielding a final set of 16,959 samples spanning 86 unique tissue and cell types (Supplementary Tables 1, 2). To enable robust learning, we required that any tissue or cell label be supported by at least two unique studies. This filtering resulted in a smaller set of 10,351 samples across 55 tissues, which, when considering ontological structure (intermediate tissue and organ system level terms), was further expanded to 72 entities (Figure 1). The remaining 6,608 samples covering 31 tissue and cell types were reserved for the label transfer evaluation. This effort represents, to our knowledge, the largest and most diverse DNAm atlas of healthy primary tissues and cell types, providing an unprecedented resource for exploration methylation patterns in normal physiology. Such a resource is critical not only for training our ontology-aware classification framework, but also for enabling future studies on tissue-specific epigenetic regulation and its implications for human health.

**Figure 1.**
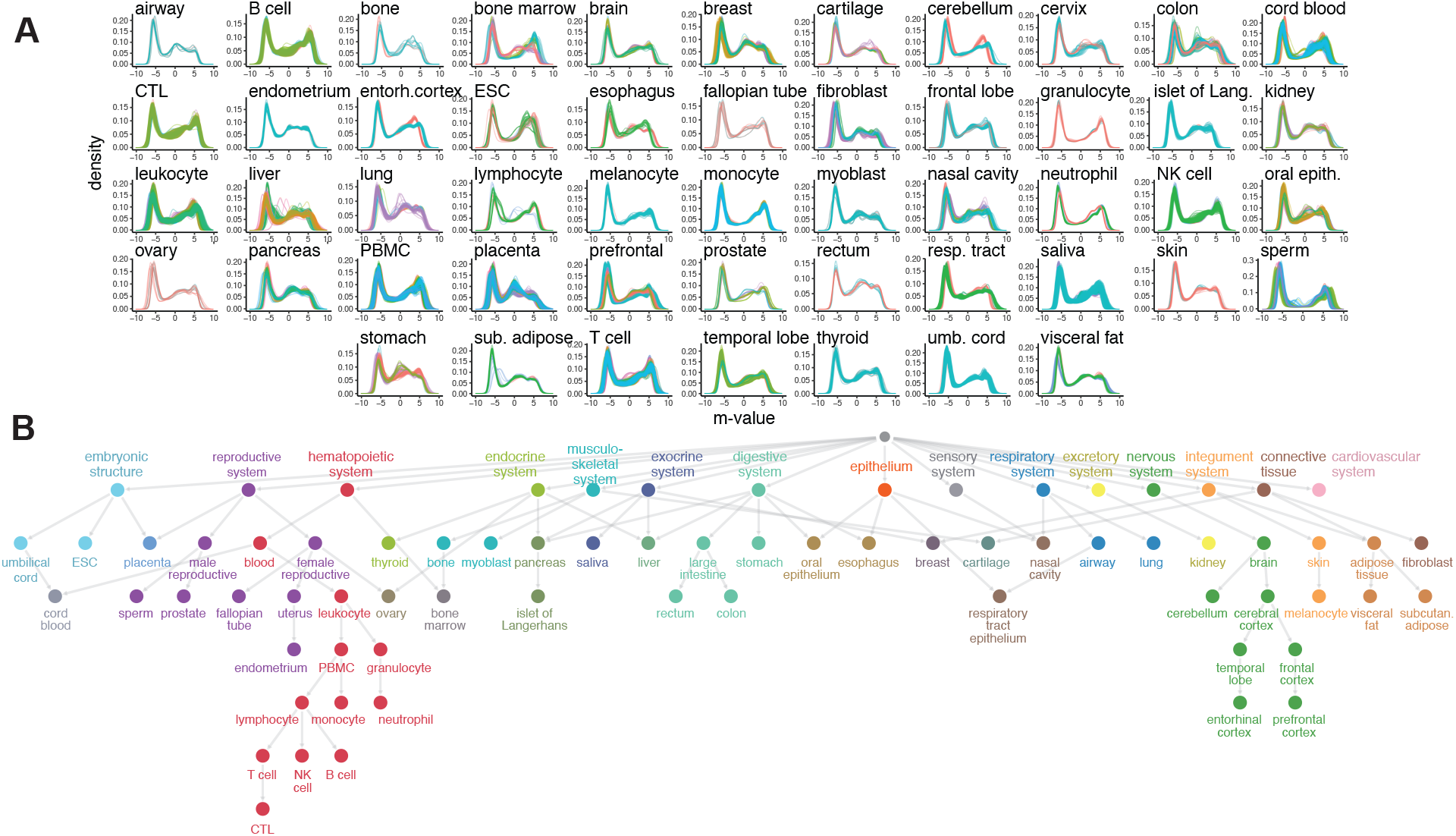
Overview of DNAm training compendium. (A) Distribution of processed M-values across all probes for all samples used for downstream learning tasks. Each line in the density plot corresponds to an individual sample, with colors indicating different datasets. (B) Anatomical ontology structure after manual curation and subsetting to tissue and cell types in the training set. Nodes are colored by their organ system, and these colors are used throughout the remainder of the paper.

To assess the consistency of our compendium, we examined M-value distributions across sample types and datasets (Figure 1A) and quantified probe variation using intraclass correlation (ICC). In general, samples of the same type exhibited similar methylation patterns (mean *ICC* = 0.97; Supplementary Figure 1, Supplementary Table 2), regardless of the number of samples per label. Sample types with greater variability likely reflect known tissue heterogeneity or cell types present in diverse bodily regions, such as the two sample types with the lowest ICCs, *bone marrow* (*ICC* = 0.92; *N* = 74) and *fibroblast* (*ICC* = 0.93; *N* = 117). Interestingly, some heterogeneous tissues like *breast* and *lung*, which both contain different mixtures of diverse cell types (e.g., fibroblasts, epithelial cells, immune cells, as well as adipocytes in the breast), still exhibited high consistency (*breast*: *ICC* = 0.98, *N* = 314; *lung*: *ICC* = 0.97, *N* = 68), suggesting that robust tissue-specific signals can be extracted without deconvolution. Notably, *neutrophils* (*ICC* = 0.99; *N* = 68) and *umbilical cord* (*ICC* = 0.99; *N* = 1, 021) had the highest internal consistency across all 450K probes.

Next, we analyzed the relationships between tissue and cell types by extracting known anatomical and functional relationships, using *is_a, part_of*, and *develops_from* annotations from the UBERON tissue ontology [26]. We combined automated extraction from UBERON with pruning and simplification of the network, as well as other functional *is_a* and *part_of* relationships from the BRENDA tissue ontology [27]. This process yielded a directed acyclic graph spanning all sample types in our compendium, organized with increasing levels of physiological resolution (Figure 1B, Supplementary Figure 2). Our tissue and cell ontology captures structural and functional heterogeneity through multiple parent nodes. For example, *breast* is a third level node with two organ system-level parents: *exocrine system* and *connective tissue*. Structurally, cell types are typically at deeper levels of the ontology, with relationships traceable back to the relevant parent organ system. For instance, the deepest node, *cytotoxic T lymphocyte (CTL)*, is functionally linked to *hematopoietic system, blood, leukocyte, PBMC, lymphocyte*, and *T cell* (Figure 1B).

**Figure 2.**
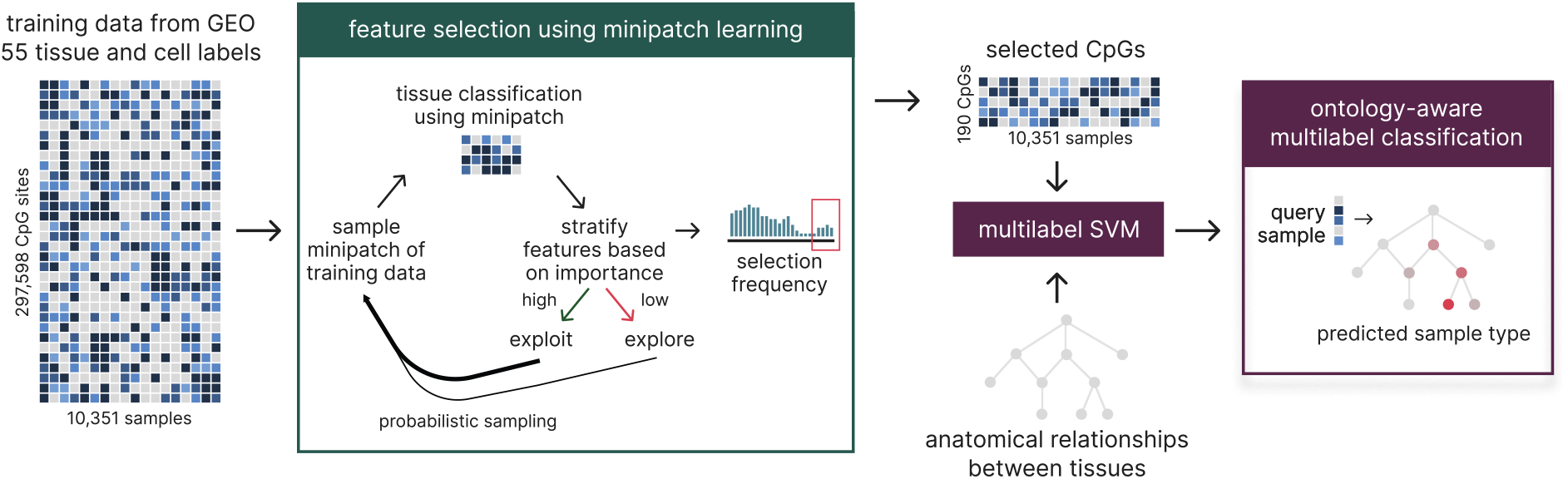
DNAm feature selection and classification workflow. We first used Minipatch learning [28] for feature selection, reducing the number of CpG features from 297,598 to 190. Selected probes are then combined with the anatomical relationships between tissues and cell types in a multi-label SVM learning framework. Given a sample of unknown origin, our ontology-aware classification framework is capable of assigning the most relevant label.

### Minipatch-based feature selection of DNAm features

To explore the functional conservation of CpGs across tissues and cell types, we devised an ontology-aware, multi-label classification framework (Figure 2). In our approach, we begin by considering the union of all quality-controlled CpG sites (297,598 sites) across the 10,351 training samples in a 3-fold cross validation setup. Importantly, cross-validation folds were grouped by study, such that samples from the same dataset were always allocated to the same fold, to ameliorate the possibility of data leakage. A key innovation of this framework is the use of Minipatch learning [28], a probabilistic sampling approach for feature selection, which iteratively refines the feature space down to a set of 190 CpG sites (Supplementary Table 3). Briefly, Minipatch learning uses decision tree-based selectors to explore different “patches” within the data by sampling small subsets of CpGs and samples, then using decision tree classifiers to evaluate their importance. The sampling probability of CpG features is increased based on their feature importance (i.e., “exploited”), while the broader feature space continues to be “explored.” After iterating until convergence, the selection frequency of a feature thus directly reflects its classification relevance. We optimized the selection frequency cutoff based on the elbow (frequency = 0.65) at which the median F1 scores on the downstream multi-label sample type classification task starts to decrease across each of the cross validation folds (fold 1 *F*_1_ = 0.80; fold 2 *F*_1_ = 0.90; fold 3 *F*_1_ = 0.87; Figure 3A). This strategy is effective and efficient, enabling sample classification approximately 200x faster compared to the traditional differential methylation approach (Figure 3B), while offering support for multi-label input. We note that the runtime difference compounds over large sample sizes, making it increasingly intractable to use a differential methylation-based classification approach on large data compendia.

**Figure 3.**
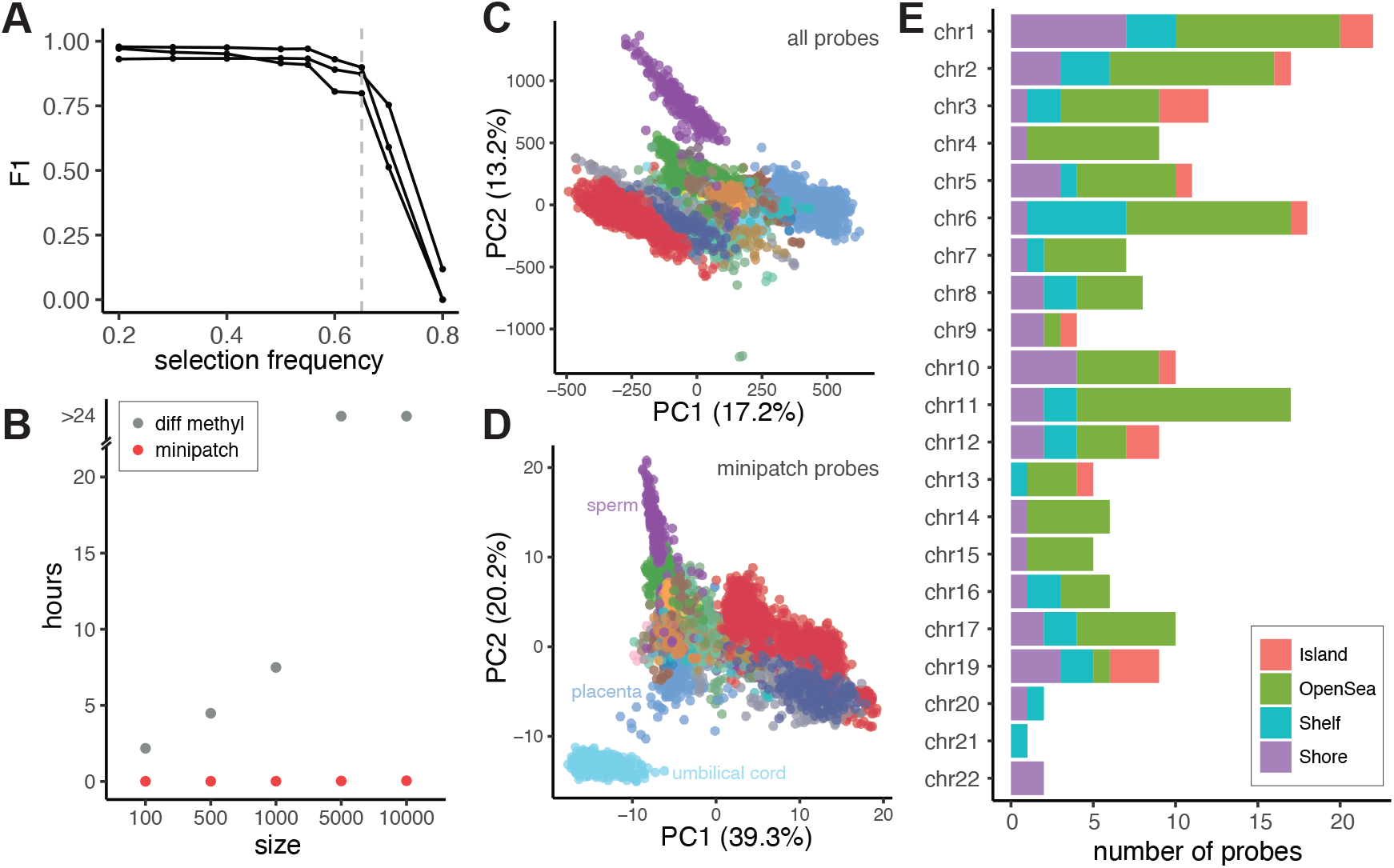
Characterization of selected CpGs for downstream classification. (A) Overall classification performance (F1 score) as a function of CpG selection frequency. Each line represents one training fold. A selection frequency of 0.65 (dotted line) was chosen to maximize performance while minimizing the total number of CpGs. (B) The runtime, in hours, as a function of the sample size of the tissue and cell type classification task when using minipatch probes versus the set of differentially methylated probes. For sample sizes 5,000 and above, the runtime for differential methylation was greater than 24 hours. (C-D) Principal component analysis of samples using the entire probe set (C) and the reduced set after Minipatch learning (D). Colors correspond to organ system as in Fig. 1B. (E) Chromosome and genomic features characterizing the 190 Minipatch-selected probes.

To further examine the CpGs selected via Minipatch learning, we visualized the DNAm feature space using principal component analysis (PCA). We found that the complete feature space of 297,598 CpGs (Figure 3C) provided some separation between sample types, but the PCA based on the 190 Minipatch-selected probes not only maintained but amplified this separation (Figure 3D). Furthermore, this reduced feature set captured substantially more variance in the first two principal components, highlighting that these probes effectively capture sample-type relevant signal.

We also characterized the genomic composition of the 190 selected CpG sites, which span all chromosomes except for chromosome 18 (Figure 3E, Supplementary Table 3). Interestingly, although the collection of probes in 450K arrays are biased towards CpG-rich regions such as CpG islands, these regions are notably underrepresented in the selected set (one-sided Fisher’s exact test, *p* = 1.9 * 10^−13^). Conversely, there is a significant over-enrichment of CpGs in open sea (one-sided Fisher’s exact test, *p* = 1.2 * 10^−6^) and shelf regions (one-sided Fisher’s exact test, *p* = 2.0 * 10^−3^), suggesting that these areas may be more useful for tissue- and cell type-specificity. We observed no significant enrichment or depletion in the shore regions (two-sided Fisher’s exact test, *p* = 0.6). This is consistent with previous reports analyzing tissue-specific differentially methylated regions, finding CpGs to be predominantly located outside of islands [29, 30] and specifically more localized to shelves and also distant regions in 450K data [18], where changes are tied with alternative transcription [31]. We also note that the relative distribution of selected CpGs per chromosome deviates from that expected by CpG site abundance alone, indicating that the selection is not simply driven by CpG site availability or chromosome length.

### Accurate cell, tissue, and organ system classification with our ontology-aware framework

To integrate ontology awareness into the framework, we used a label propagation strategy where each tissue or cell type label is defined to include not only samples explicitly annotated with that label but also those bearing more specific child labels based on the hierarchical structure of the ontology (Figure 1). For example, for the *lymphocyte* class, positive instances include samples labeled as *lymphocyte, T cell, NK cell, B cell*, or *CTL*. This allows for the training of a multi-label support vector machine (SVM) classifier that make predictions across the entire ontology, enabling nuanced classification from general organ systems to more specific tissue or cell types. In addition, each tissue or cell type prediction has an associated probability, estimated via Platt scaling, such that when there are no labels that achieve greater than 0.5 probability, the model returns ‘no prediction’, which can reduce false positives while highlighting opportunities for improved training set coverage.

Using our 3-fold cross-validation training setup, we applied this ontology-aware classification framework to all 10,351 samples and evaluated its performance (Figure 4, Supplementary Figures 3-5). As a baseline, we compared our method to a naive approach that uses tissue- and cell type–specific probes derived from differential methylation (Figure 4A, Supplementary Tables 4-5). In the naive approach, significantly differentially methylated CpGs (Holm-Sidak, *α* = 0.05) unique to each tissue are combined into a union set, which is then used to calculate sample-to-sample correlations. The tissue or cell type label with the highest average correlation is selected as the prediction. Across the 51 training labels that had sufficient direct sample annotations for differential methylation analyses, our method had significantly higher precision (one-sided Wilcoxon signed-rank test, *p* = 1.2 * 10^3^). Furthermore, our ontology-aware model was able to leverage the hierarchical relationships when direct annotations were unavailable to make predictions for an additional 21 tissues and cell type terms (average precision=0.86). Even in tissues with lower precision, our approach maintained comparable performance to the naive differential methylation method, with the exception of *NK cell*. In this case, the most frequently predicted labels were parent terms (e.g., lymphocyte, PBMC) as well as other sibling and related terms (e.g., *T cell, CTL*), indicating that misclassifications still reflected biologically relevant relationships within the ontology.

**Figure 4.**
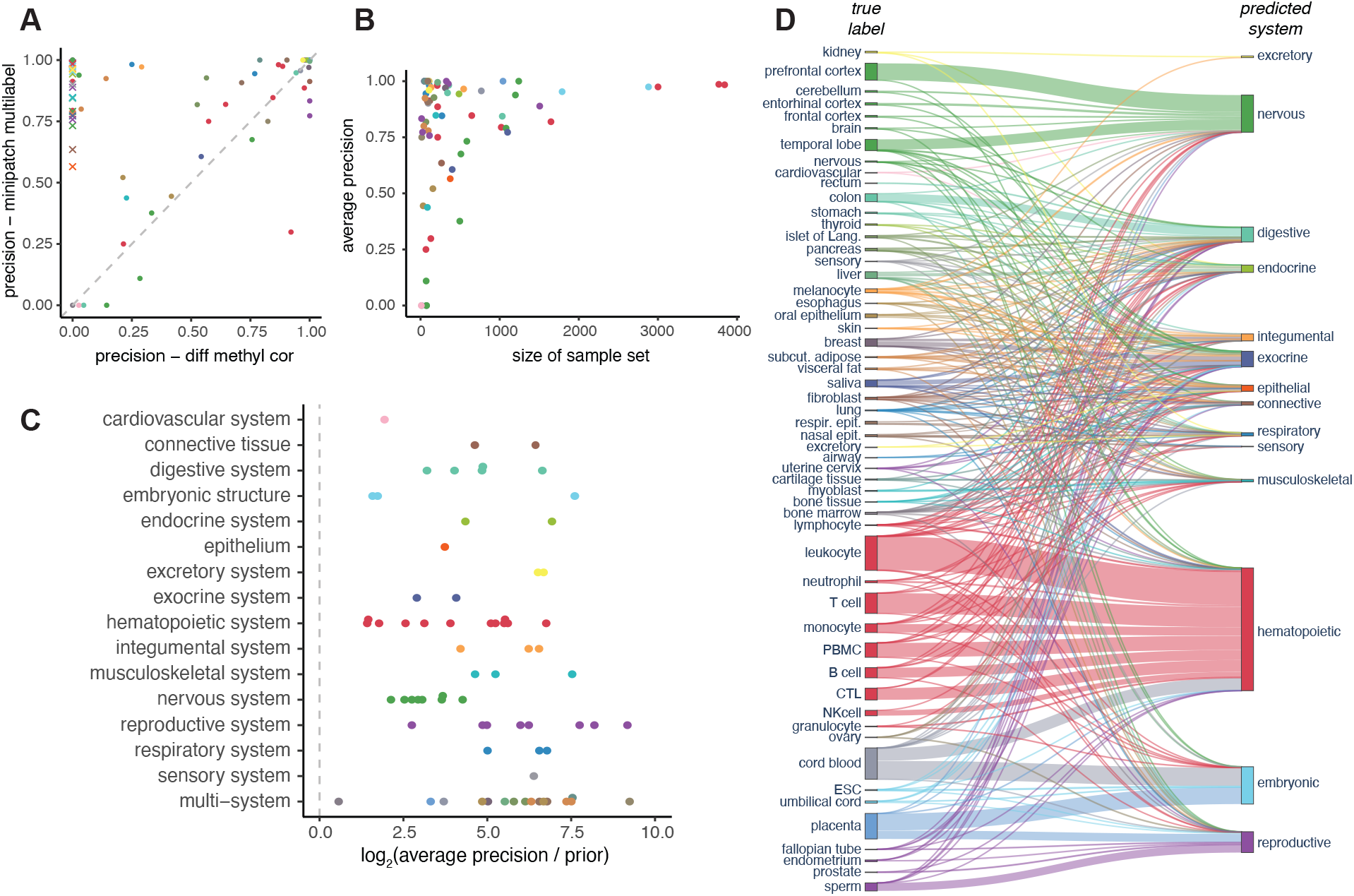
Multi-label SVM performance across tissue and cell types. (A) Average tissue-wise precision compared to a baseline method that assigns labels based on correlation with samples using probes that were significantly differential methylated. Each dot represents the average precision for a single tissue or cell type, colored by organ system. Tissues drawn as ‘x’s are intermediate nodes that cannot be predicted using differential methylation due to the lack of directly annotated labels. (B) Average tissue-wise precision as a function of the number of samples with the corresponding tissue label in the training set. Colors correspond to organ system as in Fig. 1B. (C) Log average precision over prior for each tissue and cell type label, organized by organ system. The dotted line indicates performance equal to prior. (D) Sankey plot of the actual sample label compared with the predicted system label colored according to the true organ system. The size of the Sankey ribbon corresponds to the sample size of each label.

To understand how sample size influences classification performance, we examined the average precision for each tissue label as a function of the number of samples (Figure 4B). Though many tissues and cell types achieve strong performance with relatively few numbers of samples, there is a general trend where performance increases and stabilizes at larger sample sizes, especially once the number of samples per label exceeds 1,000 (average precision=0.89). Meanwhile, all but one of the tissue or cell types with lower than average precision had fewer than 400 samples. When performance was compared against each tissue label’s priors (proportion of positive labels), to account for class label imbalance, all 72 tissues in the ontology outperformed their respective priors (Figure 4C).

To assess our model’s ability to capture broader ontological relationships, we analyzed predictions at the organ system level by comparing the predicted system nodes to true labels. Visualizing this on a Sankey diagram connecting true and predicted labels, colored according to their true systems, we see that the overwhelming majority of predictions are color-coherent, indicating that predictions either match the true label or belong to the same system (Figure 4D). This analysis also clearly highlighted multi-system nodes, where a single tissue or cell label is associated with two or more organ systems, as shown by their mixed colored ribbons in the Sankey diagram. Notable examples include *breast, nasal epithelium, placenta*, and *umbilical cord blood*; for instance, *umbilical cord blood* correctly maps to both the *hematopoietic* and *embryonic* systems, while *placenta* is also linked accurately to both the *embryonic* and *reproductive* systems.

### Ontology-aware learning enables robust predictions on unseen labels

A major advantage of our method is the ability to leverage the ontology to predict relevant, related sample type labels, even when the original sample type is absent from the training data. To evaluate this capability, we used our ontology-aware classification model to make predictions for all samples that were curated as part of our data resource but were excluded from training due to having insufficient numbers of samples or distinct studies. This resulted in a set of predictions for 6,608 samples across 31 unseen tissue and cell type labels, spanning 11 organ systems (Supplementary Table 1). Because the true target labels were not part of our training ontology, we devised an evaluation metric that measures how close each prediction is to the true target label if it were incorporated into the training ontology (Fig. 5A, Supplementary Figure 2). Specifically, we used an adjusted ontology distance metric that calculates each sample’s average graph distance from its predicted labels to the true label, accounting for the minimum distance from any training node to the target label. An optimal prediction would have a distance of 0, with larger distances indicating more disparate predictions. To account for variations in the ontology structure, we sampled 1,000 random nodes from our training ontology and measured their distances to the target node as a baseline.

**Figure 5.**
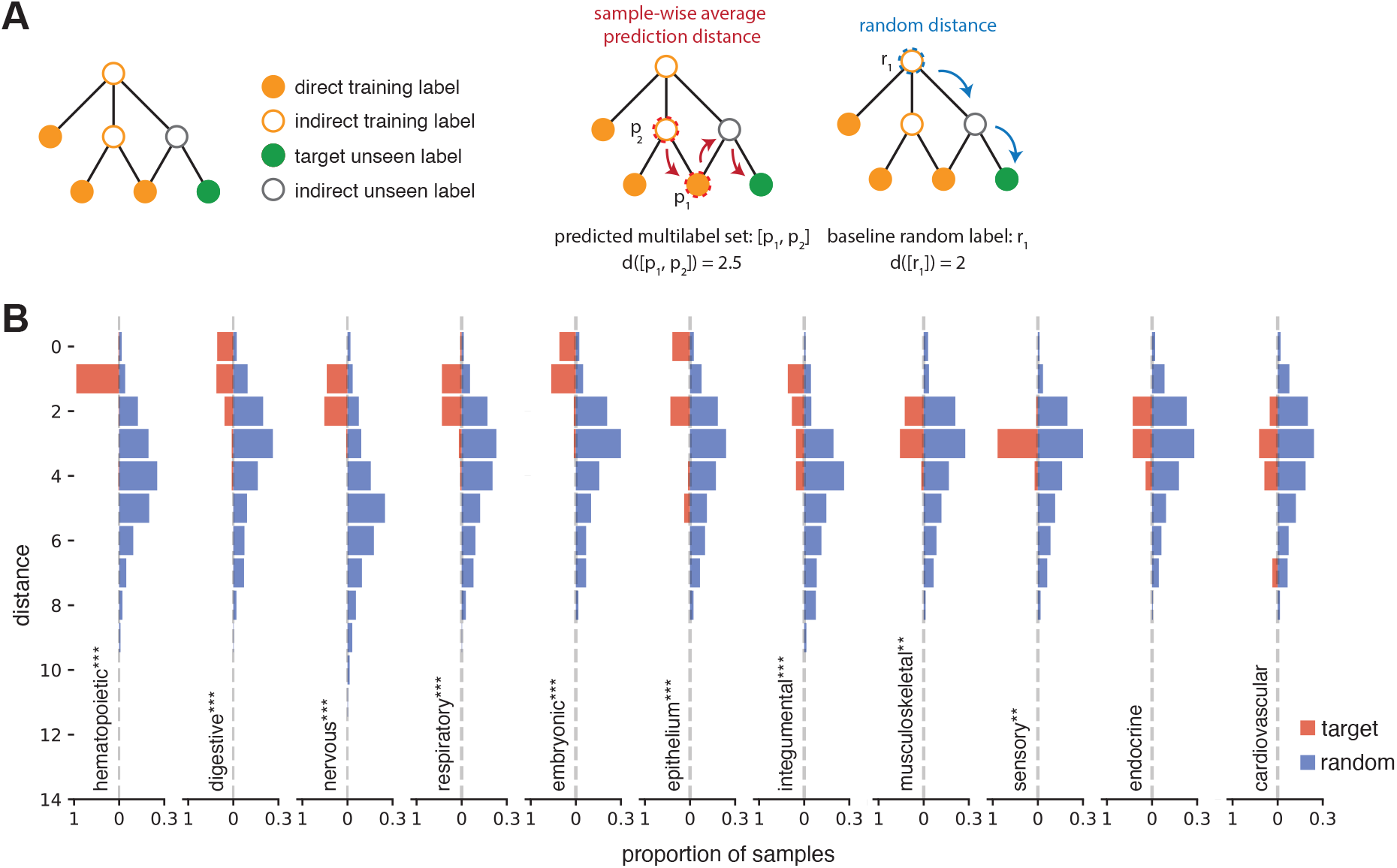
Label transfer learning evaluation. (A) Schematic of the label transfer evaluation, given the full tissue and cell ontology, including both label transfer and training set labels (Supplementary Figure 2). Given a query sample, we calculated the graph distance between every tissue label in the predicted label set (dashed red) and the target, previously unseen label (filled green circle). The final score is an average over all predicted labels for a given sample. In the random distance case, we calculated the distances for 1,000 randomly selected labels to the target label to obtain a background distribution of graph distances. (B) Histograms show the results of the label transfer evaluation, with samples grouped by organ system and ordered by significance. Sample-wise average prediction distances to the target sample are shown in red, while the background distributions of 1,000 random labels are shown in blue. All distances are adjusted such that the optimal distance when predicted correctly is equal to 0. Higher distances indicate a worse set of multi-label predictions. Asterisks represent significance compared to random using Wilcoxon rank-sum test (* *p <* 0.05, ** *p <* 0.01, *** *p <* 0.001).

Our method significantly outperforms the random baseline for 22 out of the 31 tissue and cell type labels (Supplementary Figure 6). When grouped by organ system, our approach consistently outperforms random predictions in nearly every system, with the exceptions of the *endocrine* and *cardiovascular* systems (Figure 5B). We speculate that the lower performance in the *endocrine system* may stem from its limited representation in the training set, since all endocrine tissues in our dataset are classified under multiple systems. Similarly, the *cardiovascular system* likely suffers from both having the least amount of training data overall and the fact that it contains only two multi-system tissue types in its unseen label evaluation set (*endothelial cell* and *aorta smooth muscle tissue*).

**Figure 6.**
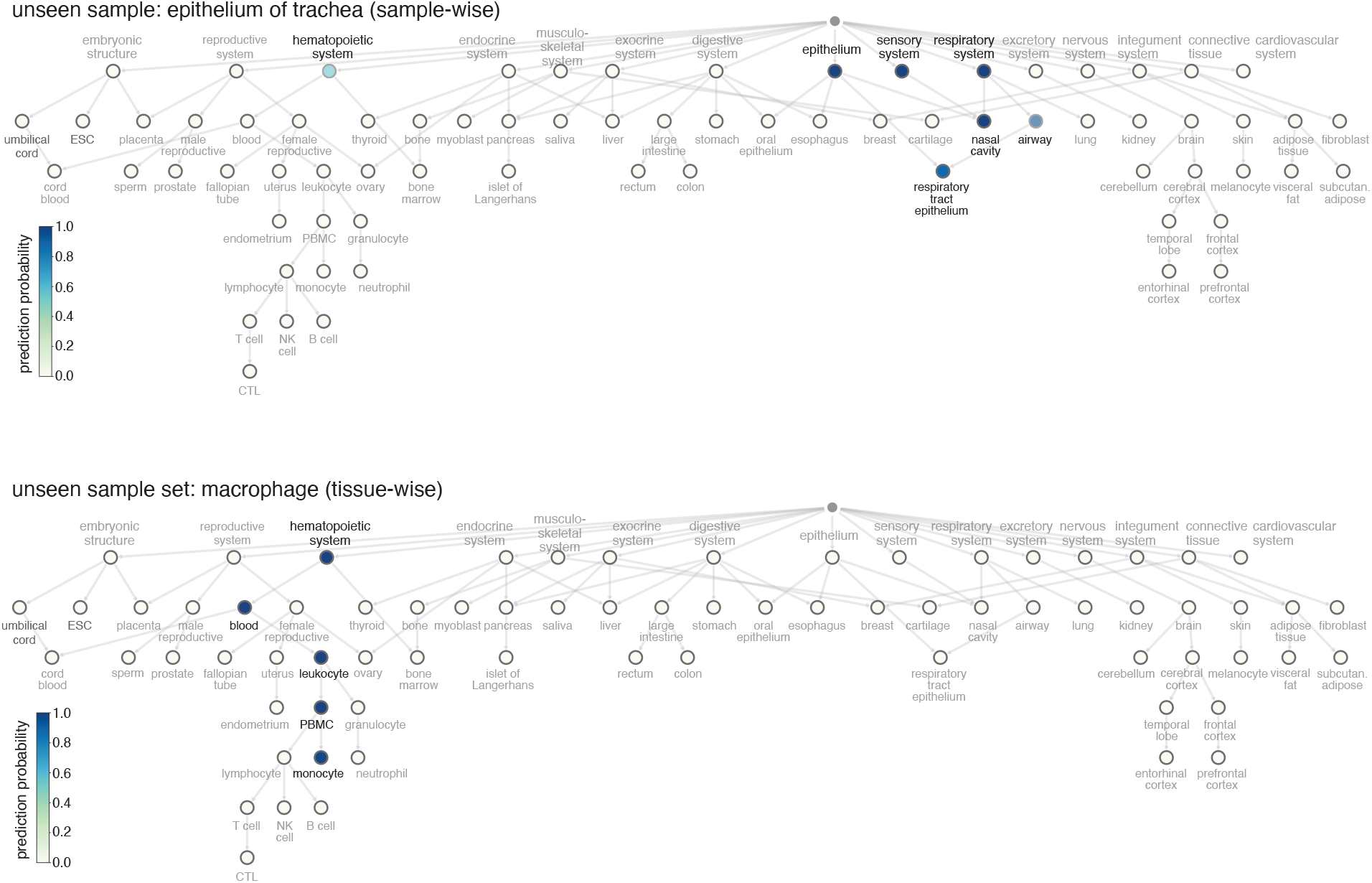
Label transfer predictions for samples with unseen labels. Example multi-label predictions are visualized on the training ontology for two different unseen cases: a single sample with the actual label of epithelium of trachea (top) and a set of samples labeled macrophage (bottom). Nodes are colored by prediction probability.

To illustrate how our ontology-driven approach can be used to interpret unseen samples, we can consider an individual sample from the *epithelium of trachea* (Figure 6 top), a tissue label unseen in our training set. Our method was able to both correctly identify its closest match, *respiratory tract epithelium* (probability = 0.87), and capture all relevant organ systems with high probability: *epithelium* (probability = 0.99), *respiratory system* (probability = 0.99), and *sensory system* (probability = 0.99), highlighting the benefit of this multi-system aware prediction framework. We can also use the same prediction scheme to consider the collection of all samples from another unseen label, *macrophage* (Figure 6 bottom). Biologically, we know that *macrophage* should be a child node of *leukocyte*. Our classifier not only captured this chain of parent relationships up through *hematopoeitic system*, but also reflected downstream functional links between monocytes and macrophages, as monocytes can differentiate into macrophages when recruited from the blood into tissues [32]. This example also highlights another use case of our method to summarize predictions across a set of samples. In general, our framework’s robust ability to place unseen labels in the context of our training data provides additional validation for the benefit of integrating structured ontological knowledge into the classification process.

## Discussion

In this study, we developed an ontology-aware multi-label classification framework leveraging DNA methylation (DNAm) data to accurately predict and characterize tissue and cell type identities. By assembling the largest curated atlas to date of exclusively healthy, primary human tissues profiled by 450K arrays, we now provide a valuable resource enabling detailed analyses of epigenetic landscapes across a diverse array of physiological systems. Importantly, our approach identifies a small subset of 190 CpGs sites that robustly distinguishes 72 tissues and cell types, demonstrating that a small number of well-selected markers can achieve high classification performance. These markers thus represent valuable reference points for establishing tissue-specific DNAm baselines, potentially aiding in the future interpretation of methylation changes associated with disease or environmental influences.

In addition, while the fact that DNAm captures lineage relationships between tissues and cell types has been well-documented [1], our findings underscore the capacity to extract and leverage functional system-related relationships in DNAm data as well. This is evidenced by the high intraclass correlation consistency across samples within heterogeneous tissues (Supplementary Figure 1, Supplementary Table 2) and the ability of our ontology-guided approach to capture both lineage and functional information, including mechanistically-related tissue and cell types in the label transfer evaluation (Figures 5 and 6, Supplementary Figure 6). Our analysis also revealed that predictive tissue- and cell-specific CpGs are predominantly localized in open sea and shelf regions rather than in CpG Islands, which corroborates previous studies of tissue- and cell-type-specific differential methylation [18, 29, 30].

Our integration of structured ontological information enabled the multi-label classifier to incorporate tissue and cell type similarity beyond explicit annotations. This setup allowed the model to make predictions at multiple levels of the ontology and to infer both direct and related labels. Beyond strong classification performance, this approach also offers practical value to biomedical researchers, facilitating more accurate annotations, improved interpretation of heterogeneously labeled datasets, and potentially uncovering novel biological insights within complex epigenetic data.

Though we present the largest DNAm data compendium of its kind, we find that sample availability and abundance remains a critical limitation, such as for the *cardiovascular* and *endocrine* systems. Currently, our ontology is limited to the sample types available from GEO measured using the Illumina Infinium HumanMethylation450 platform. Expanding data collection and curation efforts to other DNAm platforms, including emerging single-cell DNAm technologies, would help further provide a richer, multi-resolution view of the epigenetic landscape.

Looking forward, an exciting extension of our ontology-based classification framework would be integration with complementary epigenetic data modalities or sparse single-cell DNAm data, enabling fine-grained, cell-specific functional analyses in both healthy and disease contexts. As the amount of data increases, we also envision adapting this approach to leverage graph-based or network-aware machine learning techniques, allowing even richer incorporation of complex, multi-faceted tissue and cell type relationships into the classification framework. In a manner analogous to how epigenetic clocks established from healthy individuals have provided valuable baselines for biological age estimation and have yielded critical insights into aging and disease susceptibility [4, 5, 33], our comprehensive atlas of healthy tissue and cell type methylation profiles establishes a foundational reference for future tissue-based analyses. Ultimately, this resource and ontology-informed modeling approach brings a new perspective to analyses of tissue and cell type methylation and paves the way toward deeper insights into tissue-specific disease processes and epigenetic regulation.

## Methods

### Data preprocessing

#### Sample downloading and curation

We used NCBI eutils to fetch and compile metadata for 59,123 human samples deposited on the Gene Expression Omnibus using the Illumina HumanMethylation 450 BeadChip (450K; platform: GPL13534) as of October 2024. Using the title and description fields of the metadata, for each sample we manually assigned a tissue or cell type label, a disease state (healthy or diseased), and treatment status (treated or untreated). Of those, we filtered samples using the following criteria: raw idat file availability, non-diseased status, and absence of experimental perturbation such as drug treatment. We further disambiguated tissue and cell type annotations by manually assigning them to the most descriptive tissue or cell term in the UBERON ontology [26] and merging functionally and physiologically similar terms, such as *buccal mucosa* and *oral epithelium* (Supplementary Table 6).

#### DNA methylation data preprocessing and normalization

Raw Illumina 450K array data were processed into beta values and background corrected using the standard Noob (normal-exponential out-of-band) method implemented in the minfi package [34]. Preliminary sample quality control excluded 81 samples based on median intensity values across control probes, reducing the final set of samples to 16,959. To account for the differences between type 1 and type 2 probes in 450K data, beta values were normalized using beta mixture quantile dilation (BMIQ) normalization [35] from the wateRmelon package [36]. Probe-level quality control was performed using detection p-values (cutoff=0.01) [34], and probes associated with single nucleotide polymorphisms [34], cross-reactive probes [37], and those located on the sex chromosomes [34] were removed, narrowing down the total number of probes to 297,598. Finally, beta values were converted to M-values, then used for downstream classification tasks.

#### Sample partitioning for training or label transfer validation

We then partitioned all remaining annotated samples into either the training set or the label transfer validation set. To ensure robust coverage in the training set, we required each tissue or cell type label to be present in at least 2 independent studies, with a minimum of 2 samples per study, and a minimum of 6 total samples per label. Then, to ensure comprehensive coverage of physiological systems in the ontology, we selected a subset of children labels not passing the training set criteria to augment system nodes with very low training sample set sizes, including *cardiovascular system, sensory system, excretory system*, and *nervous system*, resulting in a set of 10,351 samples (Supplementary Table 1). The remaining tissue labels that did not meet the coverage criteria were used in the label transfer validation set. Samples annotated generally as *blood* without cell types were also used as part of the label transfer validation set.

#### Within tissue or cell type sample consistency evaluation

To measure the variation among directly annotated samples from the same tissue or cell type in the training set, we calculated the intraclass correlation coefficient (ICC), a statistical measure that evaluates the consistency of observations within defined groups. ICC values were calculated using the two-way consistency model based on single-measurement units implemented in the *irr* R package [38]. Higher ICC values indicate greater consistency among samples, reflecting stable tissue- or cell-type-specific methylation signatures.

#### Genomic annotation of CpG probes

To link methylation probes with genomic and transcriptional features, we mapped Illumina hg19 coordinates to hg38 using LiftOver [39]. Probes were annotated to CpG island, shelf, shore, and open sea as defined by the Illumina manifest: islands as regions with length >500 bp and >55% GC, shores as <2kb from islands, shelves as <2kb from shores, and the remaining as open sea [40].

#### Tissue and cell ontology

In order to systematically define the physiological relationships between tissue and cell type labels, we leveraged the extended UBERON ontology [26], which includes the Cell Ontology, together with the BRENDA tissue ontology [27], which captures functional anatomical tissue relationships. We manually curated each ontology edge for all tissue and cell type labels in our complete data compendium (including both training and label transfer sets) as well as indirectly connected entities either through neighboring or intermediate connections between observed sample annotations. Specifically, the edge curation was based on existing *is_a, part_of*, and *develops_from* relationships in the UBERON and BRENDA ontologies to build a biologically intuitive directed acyclic graph (DAG) primarily organized by organ system. This resulted in a DAG with 118 nodes and 139 edges including the root node. To obtain the smaller training ontology, we subsetted entries of the ontology to contain only entities for tissues or cell types in the training set or associated indirect nodes, resulting in a training DAG with 72 entities joined by a root node and 88 edges.

#### CpG feature selection using Minipatch learning

For feature selection, we used the Minipatch learning method [41]. Briefly, Minipatch learning iteratively selects random subsets of features and samples, referred to as ‘minipatches’ and assesses feature importance using decision trees for multiclass tissue and cell type classification. The resulting feature importance from each patch are then combined with cumulative importance values, which then inform feature selection probabilities for subsequent minipatches. Through this iterative sampling approach, each feature’s utility across the entire dataset can be efficiently estimated based on its overall selection frequency, enabling identification of a small set of highly informative CpGs.

#### Hyperparameter optimization and cross-validation

Hyperparameters for feature selection and classification were set to default values, with the exceptions of minipatch size and selection frequency threshold. Recommended Minipatch learning parameters for the size of sample and feature ratios per minipatch were used 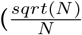, where *N* is the total number of samples or features, respectively) [28]. Selection frequency threshold was optimized using three-fold cross-validation implemented with scikit-learn [42]. Cross-validation folds were stratified by tissue labels and grouped by dataset, ensuring that samples from a single dataset were restricted to the same fold to avoid data leakage. Within each fold, we assessed Minipatch learning and classification performance using F1 score as the evaluation metric. We determined the selection frequency threshold by identifying the elbow point of classification performance across cross-validation folds (selection frequency=0.65), and a final Minipatch learning and multi-label classifier with optimized hyperparameters on the entire training dataset were used for downstream predictions.

#### Ontology-aware multi-label classification

To effectively leverage hierarchical relationships among tissues and cell types present in our ontology, we propagated nodes labels through the DAG, such that each sample’s annotation includes not only its directly annotated label, but also any parent labels. Thus, our classifier could learn methylation patterns associated with both precise tissue and cell types as well as more general signals, including for organ system level nodes. For classification, we used a multi-label support vector machine (SVM) with balanced class weights and a linear kernel, implemented via scikit-learn [42]. The probabilities were calculated using Platt scaling [42], and those with over 0.5 predicted probabilities were considered as positive predictions. In situations where no labels had predicted probabilities greater than 0.5, it was considered as ‘no prediction.’

We define *n* as the number of samples, indexed by *i*, with true and predicted label sets **y**_*i*_ and **ŷ**_*i*_, respectively. Sample-wise accuracy (Eq. 1) measured exact matches, while the Jaccard index (Eq. 2) quantified partial overlap using *TP*_*i*_, *FP*_*i*_, and *FN*_*i*_. Sample-wise precision (Eq. 3) averaged per-sample scores, while tissue-wise precision (Eq. 4) and F1-score (Eq. 5) were computed per tissue and summarized by the median. Tissue-wise metrics were only computed for tissues with at least one positive sample.

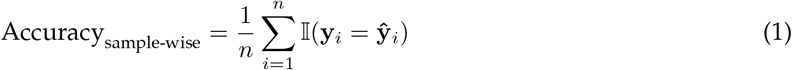

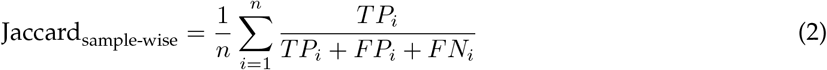

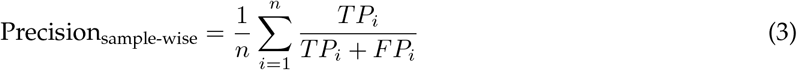

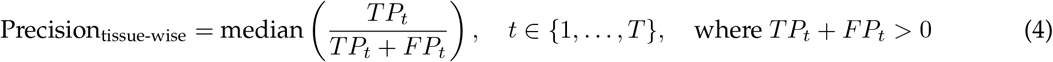

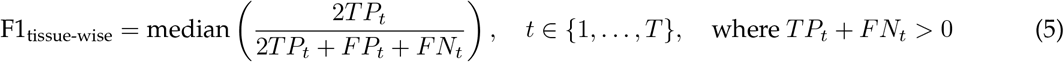

#### Differential methylation baseline

Differential methylation was used as a baseline feature selection method due to its widespread use as a standard analytical approach in DNA methylation studies, particularly for identifying tissue-specific methylation markers. We calculated tissue-specific differentially methylated probes via one-versus-rest comparisons while accounting for dataset labels as covariates using the Python package Methylize [43]. Probes were considered significantly differentially methylated if they passed an alpha of 0.05 after multiple hypothesis test correction using the Statsmodels package [44]. For classification based on differential methylation, we used the union set of differentially methylated probes across all tissues. Within the established cross-validation folds, we calculated sample-to-sample Pearson correlation coefficients between samples in training folds with those in the validation fold. Tissue and cell type predictions for each sample in the validation fold were assigned based on the tissue or cell type label corresponding to the highest average correlation coefficient of grouped training samples. The metrics reported are averages across all cross-validation folds.

#### Runtime evaluations

Runtime evaluations for both differential methylation and Minipatch learning for feature selection were restricted to 10 threads on a server with 4 Intel(R) Xeon(R) Gold 5220 CPUs and 1.5TB RAM. Only library-internal parallelization was permitted, thus differential methylation was performed using parallelization and minipatch learning was not. For evaluation purposes, we created subsampled datasets with 100, 500, 1,000, 5,000, and 10,000 samples from the entire training set, stratifying by tissue labels to maintain consistent representation of tissue labels across runtime measurements. Measured runtimes include only the time required to fit each feature selection method and not downstream classifier training and prediction.

#### Label transfer evaluation

To assess prediction performance in the label transfer evaluation, we devised a graph distance-based metric for each sample that measures the average distances of each predicted label generated by the multi-label classifier to the annotated true label. Specifically, we used an undirected version of our ontology that included both the training and unseen labels (Supplementary Figure 2). Because predictions would be assigned to labels present in the training set, we computed an adjusted ontology distance that took into account the nearest available training node for each unseen label. More formally, let *d*_*i*_ represent the ontology distance between the target (unseen) label and the *i*-th predicted label for a given sample. We first identified the minimum distance between the target node and all *n* nodes in the training set:

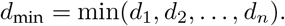

We then computed the adjusted ontology distance by subtracting this minimum achievable distance from each prediction’s ontology distance:

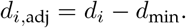

Thus, an adjusted distance of 0 indicates an optimal prediction (matching the closest possible training node), while adjusted distances reflect less accurate label transfer.

To contextualize our label transfer performance, we established random baseline distributions for each unseen label. This involved randomly sampling labels with replacement from the full set of labels in the training ontology (1,000 iterations per unseen label). For each sampled label, we calculated the adjusted ontology distance to the target unseen label using the same method described above.

## Data and code availability

The full data compendium (M-values from all 16,959 downloaded, preprocessed, normalized samples), annotations (Supplementary Table 1), curated ontology, and all analysis code can be accessed via github https://github.com/ylaboratory/methylation-classification.

## Declaration of interests

The authors declare no competing interests.

## Acknowledgments

The authors would like to thank members of the ylaboratory for helpful discussions. This work was supported by the Cancer Prevention & Research Institute of Texas [CPRIT RR190065 to VY] and the National Science Foundation [NSF DBI-2144534 to VY and NSF DMS-1554821 to GA]. VY is a CPRIT Scholar in Cancer Research.

## Notes

### Competing Interest Statement

The authors have declared no competing interest.

## References

1. Dor, Y. & Cedar, H. Principles of DNA methylation and their implications for biology and medicine. The Lancet 392, 777–786 (2018).

2. Ziller, M. J. et al. Charting a dynamic DNA methylation landscape of the human genome. Nature 500, 477–481 (2013).

3. Loyfer, N. et al. A DNA methylation atlas of normal human cell types. Nature 613, 355–364 (2023).

4. Horvath, S. DNA methylation age of human tissues and cell types. en. Genome Biology 14, R115. ISSN: 1465-6906. http://genomebiology.biomedcentral.com/articles/10.1186/gb-2013-14-10-r115 (2024) (2013).

5. Hannum, G. et al. Genome-wide Methylation Profiles Reveal Quantitative Views of Human Aging Rates. en. Molecular Cell 49, 359–367. ISSN: 10972765. https://linkinghub.elsevier.com/retrieve/pii/S1097276512008933 (2024) (Jan. 2013).

6. Sokolov, A. V. & Schiöth, H. B. Decoding depression: a comprehensive multi-cohort exploration of blood DNA methylation using machine learning and deep learning approaches. Translational Psychiatry 14, 287 (2024).

7. Yang, Z. et al. Correlation of an epigenetic mitotic clock with cancer risk. Genome biology 17, 1–18 (2016).

8. Zheng, S. C., Breeze, C. E., Beck, S. & Teschendorff, A. E. Identification of differentially methylated cell types in epigenome-wide association studies. Nature methods 15, 1059–1066 (2018).

9. Teschendorff, A. E., Breeze, C. E., Zheng, S. C. & Beck, S. A comparison of reference-based algorithms for correcting cell-type heterogeneity in Epigenome-Wide Association Studies. BMC bioinformatics 18, 1–14 (2017).

10. Titus, A. J., Gallimore, R. M., Salas, L. A. & Christensen, B. C. Cell-type deconvolution from DNA methylation: a review of recent applications. Human molecular genetics 26, R216–R224 (2017).

11. Yousefi, P. D. et al. DNA methylation-based predictors of health: applications and statistical considerations. Nature Reviews Genetics 23, 369–383 (2022).

12. Ji, H. et al. Comprehensive methylome map of lineage commitment from haematopoietic progenitors. en. Nature 467, 338–342. ISSN: 0028-0836, 1476-4687. https://www.nature.com/articles/nature09367 (2025) (Sept. 2010).

13. Teschendorff, A. E. & Relton, C. L. Statistical and integrative system-level analysis of DNA methylation data. Nature Reviews Genetics 19, 129–147 (2018).

14. Luo, C. et al. Single-cell methylomes identify neuronal subtypes and regulatory elements in mammalian cortex. en. Science 357, 600–604. ISSN: 0036-8075, 1095-9203. https://www.science.org/doi/10.1126/science.aan3351 (2025) (Aug. 2017).

15. Mulqueen, R. M. et al. Highly scalable generation of DNA methylation profiles in single cells. en. Nature Biotechnology 36, 428–431. ISSN: 1087-0156, 1546-1696. https://www.nature.com/articles/nbt.4112 (2025) (May 2018).

16. Iqbal, W. & Zhou, W. Computational Methods for Single-Cell DNA Methylome Analysis. en. Genomics, Proteomics & Bioinformatics 21, 48–66. ISSN: 1672-0229, 2210-3244. https://academic.oup.com/gpb/article/21/1/48/7274167 (2025) (Feb. 2023).

17. Ahn, J., Heo, S., Lee, J. & Bang, D. Introduction to Single-Cell DNA Methylation Profiling Methods. en. Biomolecules 11, 1013. ISSN: 2218-273X. https://www.mdpi.com/2218-273X/11/7/1013 (2025) (July 2021).

18. Lokk, K. et al. DNA methylome profiling of human tissues identifies global and tissue-specific methylation patterns. Genome biology 15, 1–14 (2014).

19. Teschendorff, A. E., Zhu, T., Breeze, C. E. & Beck, S. EPISCORE: cell type deconvolution of bulk tissue DNA methylomes from single-cell RNA-Seq data. Genome biology 21, 1–33 (2020).

20. Moss, J. et al. Comprehensive human cell-type methylation atlas reveals origins of circulating cell-free DNA in health and disease. Nature communications 9, 5068 (2018).

21. Li, S. et al. Comprehensive tissue deconvolution of cell-free DNA by deep learning for disease diagnosis and monitoring. Proceedings of the National Academy of Sciences 120, e2305236120 (2023).

22. Varley, K. E. et al. Dynamic DNA methylation across diverse human cell lines and tissues. Genome research 23, 555–567 (2013).

23. Ziller, M. J. et al. Charting a dynamic DNA methylation landscape of the human genome. en. Nature 500, 477–481. ISSN: 0028-0836, 1476-4687. http://www.nature.com/articles/nature12433 (2022) (Aug. 2013).

24. Edgar, R. Gene Expression Omnibus: NCBI gene expression and hybridization array data repository. Nucleic Acids Research 30, 207–210. ISSN: 13624962. https://academic.oup.com/nar/article-lookup/doi/10.1093/nar/30.1.207 (2022) (Jan. 2002).

25. Bibikova, M. et al. High density DNA methylation array with single CpG site resolution. Genomics 98, 288–295 (2011).

26. Mungall, C. J., Torniai, C., Gkoutos, G. V., Lewis, S. E. & Haendel, M. A. Uberon, an integrative multi-species anatomy ontology. en. Genome Biology 13, R5. ISSN: 1465-6906. http://genomebiology.biomedcentral.com/articles/10.1186/gb-2012-13-1-r5 (2024) (2012).

27. Chang, A. et al. BRENDA, the ELIXIR core data resource in 2021: new developments and updates. en. Nucleic Acids Research 49, D498–D508. ISSN: 0305-1048, 1362-4962. https://academic.oup.com/nar/article/49/D1/D498/5992283 (2025) (Jan. 2021).

28. Yao, T. & Allen, G. I. Feature Selection for Huge Data via Minipatch Learning arXiv:2010.08529 [cs, stat]. Feb. 2021. http://arxiv.org/abs/2010.08529 (2024).

29. Doi, A. et al. Differential methylation of tissue- and cancer-specific CpG island shores distinguishes human induced pluripotent stem cells, embryonic stem cells and fibroblasts. en. Nature Genetics 41, 1350–1353. ISSN: 1061-4036, 1546-1718. https://www.nature.com/articles/ng.471 (2025) (Dec. 2009).

30. Irizarry, R. A. et al. The human colon cancer methylome shows similar hypo- and hypermethylation at conserved tissue-specific CpG island shores. en. Nature Genetics 41, 178–186. ISSN: 1061-4036, 1546-1718. https://www.nature.com/articles/ng.298 (2025) (Feb. 2009).

31. Slieker, R. C. et al. Identification and systematic annotation of tissue-specific differentially methylated regions using the Illumina 450k array. en. Epigenetics & Chromatin 6, 26. ISSN: 1756-8935. https://epigeneticsandchromatin.biomedcentral.com/articles/10.1186/1756-8935-6-26 (2025) (Dec. 2013).

32. Yang, J., Zhang, L., Yu, C., Yang, X.-F. & Wang, H. Monocyte and macrophage differentiation: circulation inflammatory monocyte as biomarker for inflammatory diseases. en. Biomarker Research 2, 1. ISSN: 2050-7771. https://biomarkerres.biomedcentral.com/articles/10.1186/2050-7771-2-1 (2025) (Dec. 2014).

33. Levine, M. E. et al. An epigenetic biomarker of aging for lifespan and healthspan. en. Aging 10, 573–591. ISSN: 1945-4589. https://www.aging-us.com/lookup/doi/10.18632/aging.101414 (2025) (Apr. 2018).

34. Aryee, M. J. et al. Minfi: a flexible and comprehensive Bioconductor package for the analysis of Infinium DNA methylation microarrays. en. Bioinformatics 30, 1363–1369. ISSN: 1367-4811, 1367-4803. https://academic.oup.com/bioinformatics/article/30/10/1363/267584 (2024) (May 2014).

35. Teschendorff, A. E. et al. A beta-mixture quantile normalization method for correcting probe design bias in Illumina Infinium 450 k DNA methylation data. en. Bioinformatics 29, 189–196. ISSN: 1367-4811, 1367-4803. https://academic.oup.com/bioinformatics/article/29/2/189/204142 (2024) (Jan. 2013).

36. Pidsley, R. et al. A data-driven approach to preprocessing Illumina 450K methylation array data. en. BMC Genomics 14, 293. ISSN: 1471-2164. https://bmcgenomics.biomedcentral.com/articles/10.1186/1471-2164-14-293 (2024) (Dec. 2013).

37. Chen, Y.-a. et al. Discovery of cross-reactive probes and polymorphic CpGs in the Illumina Infinium HumanMethylation450 microarray. en. Epigenetics 8, 203–209. ISSN: 1559-2294, 1559-2308. http://www.tandfonline.com/doi/abs/10.4161/epi.23470 (2025) (Feb. 2013).

38. Gamer, M., Lemon, J. &, I. F. P. S. irr: Various Coefficients of Interrater Reliability and Agreement R package version 0.84.1 (2019). https://CRAN.R-project.org/package=irr.

39. Hinrichs, A. S. The UCSC Genome Browser Database: update 2006. en. Nucleic Acids Research 34, D590–D598. ISSN: 0305-1048, 1362-4962. https://academic.oup.com/nar/article-lookup/doi/10.1093/nar/gkj144 (2025) (Jan. 2006).

40. Zhou, W., Laird, P. W. & Shen, H. Comprehensive characterization, annotation and innovative use of Infinium DNA methylation BeadChip probes. en. Nucleic Acids Research, gkw967. ISSN: 0305-1048, 1362-4962. https://academic.oup.com/nar/article-lookup/doi/10.1093/nar/gkw967 (2025) (Oct. 2016).

41. Yao, T. & Allen, G. I. Feature Selection for Huge Data via Minipatch Learning. Publisher: arXiv Version Number: 2. https://arxiv.org/abs/2010.08529 (2023) (2020).

42. Pedregosa, F. et al. Scikit-learn: Machine Learning in Python. Journal of Machine Learning Research 12, 2825–2830 (2011).

43. FOXO Bioscience. methylize https://life-epigenetics-methylprep.readthedocs-hosted.com/en/latest/index.html.

44. Seabold, S. & Perktold, J. statsmodels: Econometric and statistical modeling with python in 9th Python in Science Conference (2010).

