## Supplementary Figures for "Ontology-aware DNA methylation classification with a curated atlas of human tissues and cell types"

### Methylation classification: supplementary materials

#### Supplementary tables

**Supplementary Table 1:** Sample information for the training and label transfer datasets.

**Supplementary Table 2:** Summarized training dataset by tissue-wise counts and ICC. ICC values were calculated only for directly annotated tissues without label propagation or supplementary samples for system-level labels.

**Supplementary Table 3:** Minipatch learning selected probes, their genomic locations, and their CpG island region information, also includes the numbers of minipatch and total probes per chromosome.

**Supplementary Table 4:** Performance comparisons for each tissue and cell type using probes selected using Minipatch learning or differential methylation (both precision and accuracy). Some tissue labels (such as system-level nodes) had insufficient direct annotations for differential methylation analysis and thus downstream performance could not be evaluated. For labels where differential methylation could be assessed, the average number of probes (across 3 folds of training) is also reported.

**Supplementary Table 5:** Tissue names and IDs merged due to physiological and functional similarities. Tissue names and IDs added to training set to aid in system-level node training and cross-validation.

#### Supplementary figures

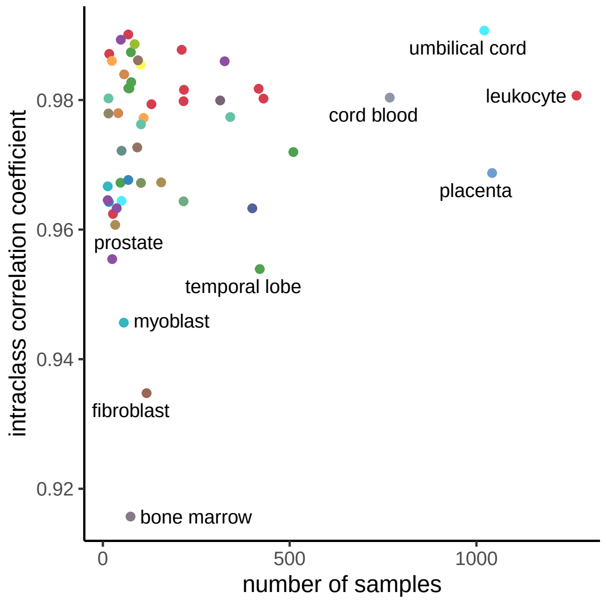


**Supplementary Figure 1:** Intraclass correlation coefficient (ICC) for each tissue label by number of samples. Tissue colors were determined according to their system-level node.

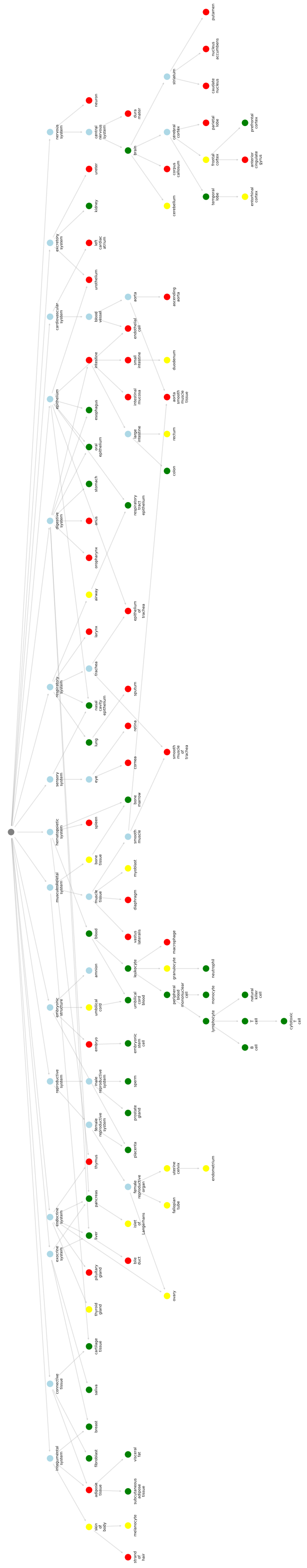

**Supplementary Figure 2:** Full ontology. Node color indicates number of studies: green (3+), yellow (2), red (1), light blue (inferred via propagation), and grey (root).

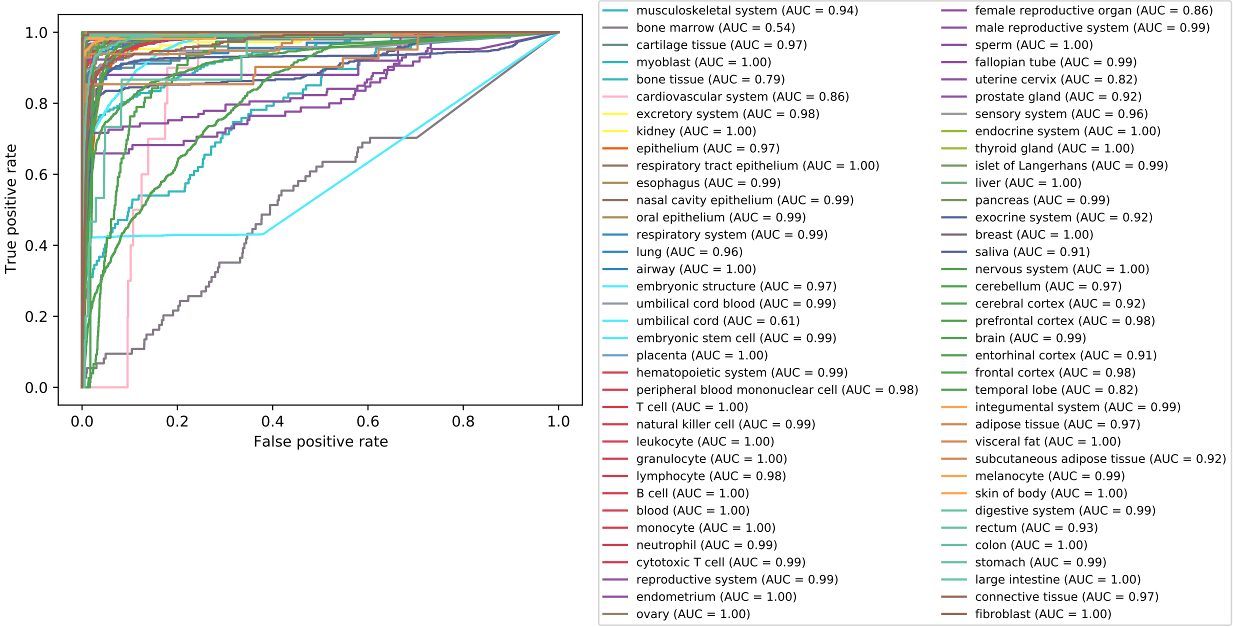


**Supplementary Figure 3:** Receiver operating characteristics and area under the curve (AUC) for each tissue colored by respective systems.

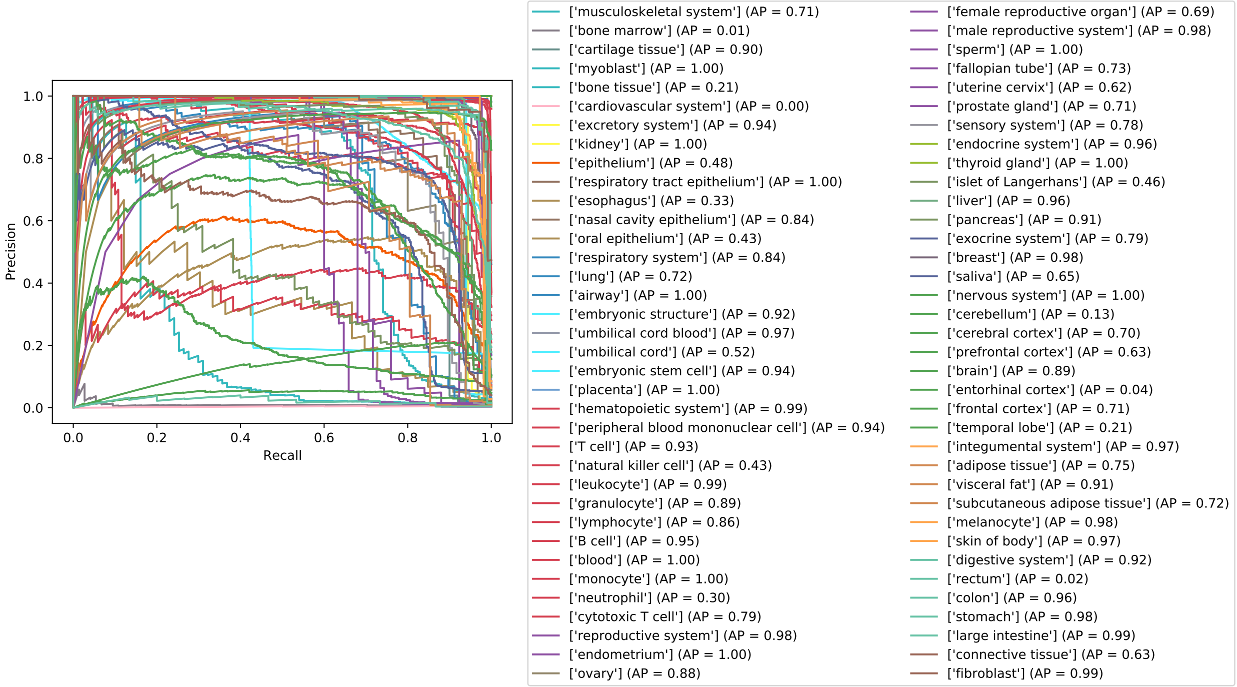


**Supplementary Figure 4:** Precision recall curve and average precision (AP) for each tissue colored by respective systems.

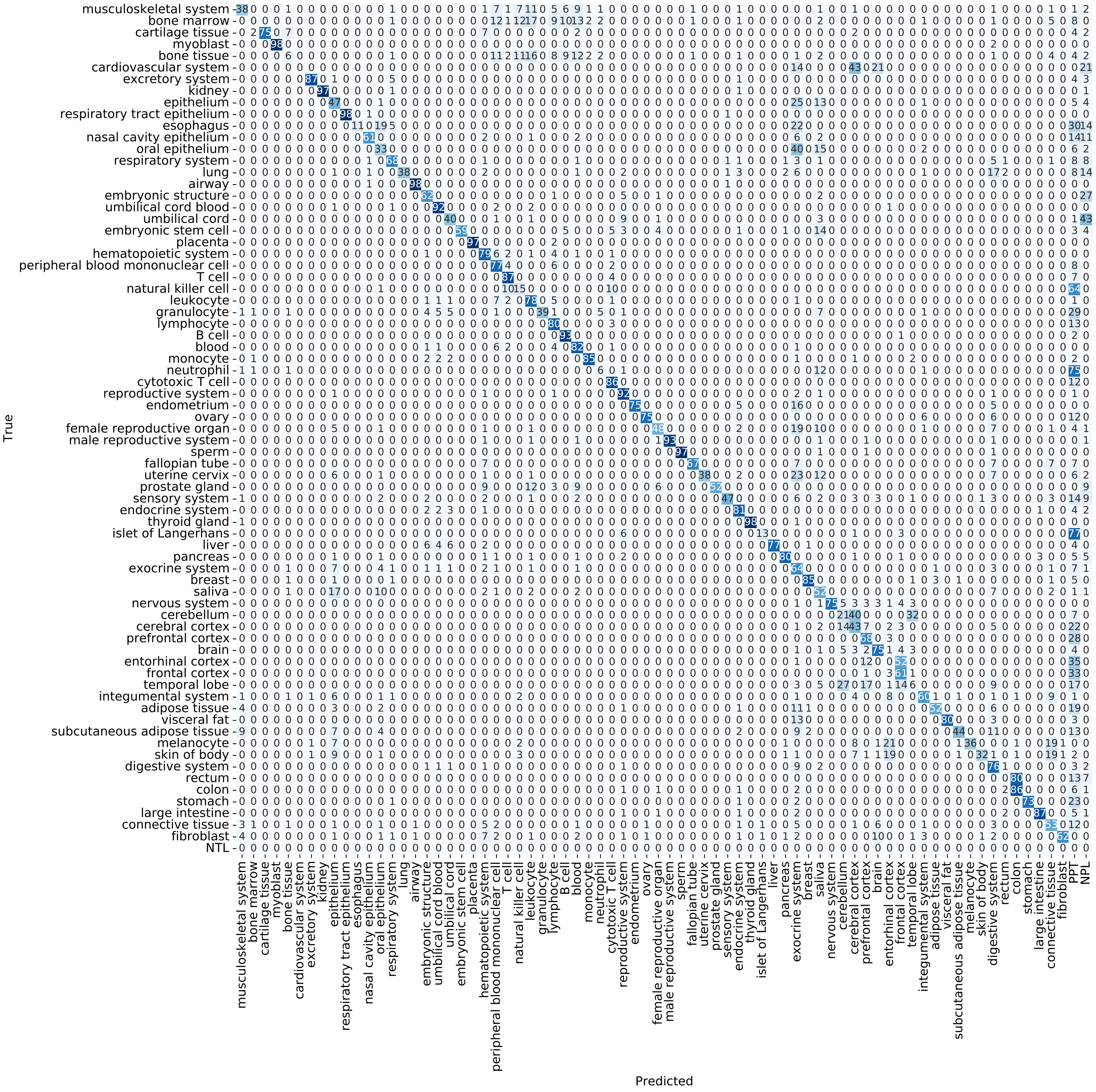


**Supplementary Figure 5:** Normalized multi-label confusion matrix for all validation folds adapted from [1]. PPT: Predicted parent term, NPL: no positive label, NTL: no true label.

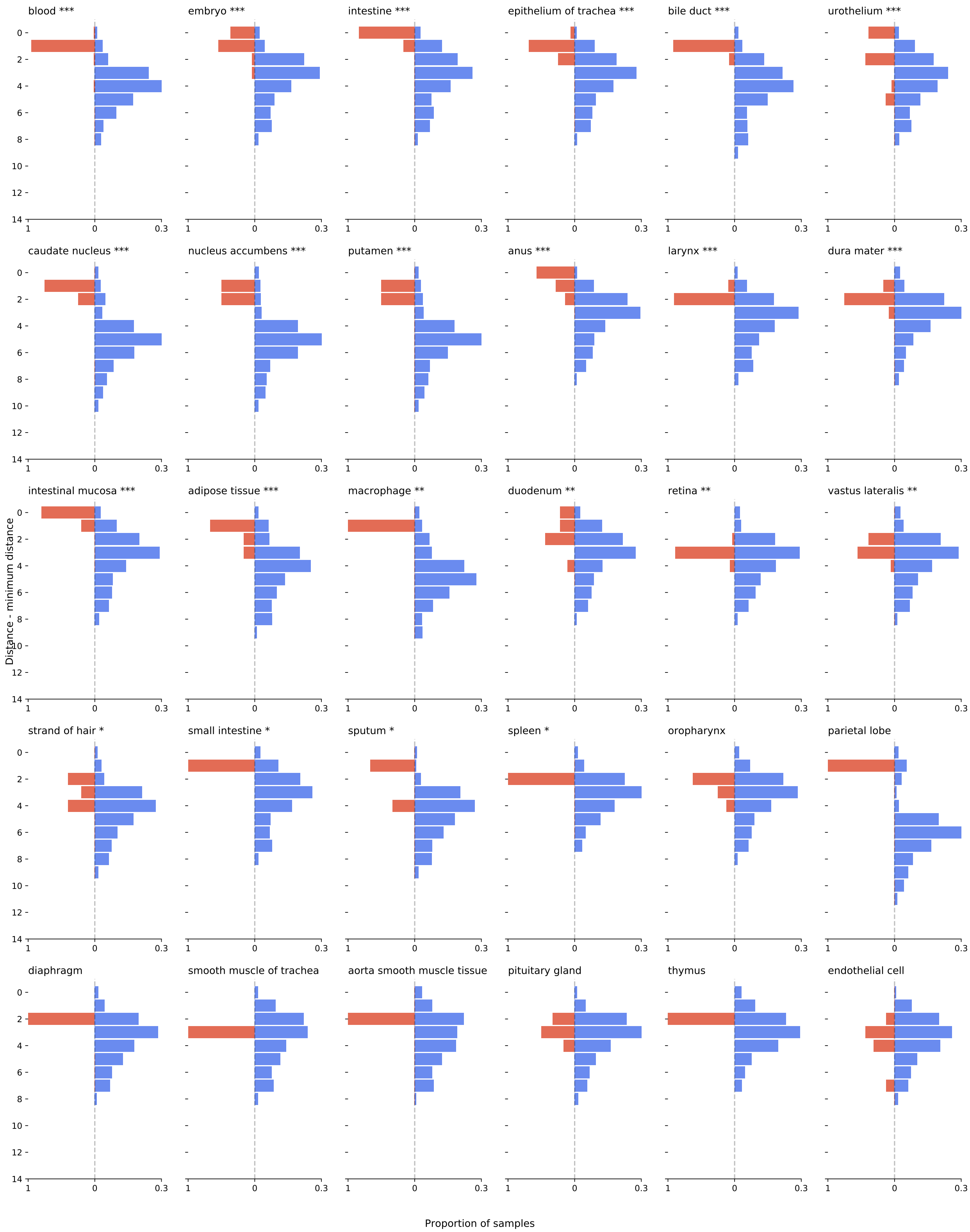


**Supplementary Figure 6:** Label transfer distances for each tissue. *:p-value<0.05, **:p-value<0.01, ***:p-value<0.001.

#### References

1. Heydarian, M., Doyle, T. E. & Samavi, R. MLCM: Multi-Label Confusion Matrix. *IEEE Access* **10,** 19083–19095. ISSN: 2169-3536. `https://ieeexplore.ieee.org/document/9711932/` (2025) (2022).
